# Spatial variation in benthic community composition on a minimally disturbed coral reef in the years following a prolonged marine heatwave

**DOI:** 10.1101/2025.02.04.636502

**Authors:** Rebecca L. G. Hansen, Andrew Halford, Dominique G. Maucieri, Julia K. Baum

## Abstract

With the increased frequency of marine heatwaves and related coral mass mortality events, it is imperative that we improve our understanding of coral reef recovery processes including how benthic community compositions can change over time. Coral reef benthic communities are influenced by many abiotic factors, but on most reefs local anthropogenic disturbances overshadow these factors thus obscuring their influence. Here, we leverage a dataset from a coral reef with very minimal local anthropogenic disturbance – the uninhabited southern coast of the world’s largest atoll (Kiritimati) – to assess spatial variation in benthic community composition three years after the mass coral mortality event driven by the 2015-2016 El Niño. Across forereef sites ranging from 7 to 22 m, scleractinian coral cover remained very low (6.9 ± 0.4 % SE) while soft coral cover was < 1%. Coral cover was highest at deep sites (18-22 m compared to sites at 7-10 m depth) and exposed locations, where stress-tolerant corals likely made up a larger proportion of the coral community before the mass mortality event. Higher cover of crustose coralline algae at shallow exposed sites, fleshy macroalgae at deep sites, and turf algae at exposed locations were consistent with taxa-specific preferences for light, wave action, and sedimentation. The abundances of the most common genera of juvenile coral (*Acropora* and *Pocillopora*) were still low (< 1 juvenile colony per genus per site) and varied only with depth. These findings demonstrate how variation in pre-mortality coral composition can lead to differences in post-mortality benthic communities on low disturbance coral reefs.

## Introduction

Climate change has increased the frequency and intensity of marine heatwaves, making mass coral bleaching events more common and severe (Frölicher et al. 2018; Oliver et al. 2018; Fox-Kemper et al. 2021; IPCC 2022). Long periods of bleaching, such as those triggered during severe marine heatwaves, can weaken corals and lead to mass mortality (Baker et al. 2008; LaJeunesse et al. 2018). As marine heatwaves are predicted to increase in the future (Oliver et al. 2018), repeat coral mass mortality events are becoming the reality for tropical reef communities, which depend on corals for structure and habitat (Jackson 1991; Hughes et al. 2018; Eakin et al. 2019). Losses in coral cover can lead to increased abundance of other benthic species such as non-coral invertebrates and macroalgae, causing both short and long-term changes in reef benthic composition (McClanahan 2000; Golbuu et al. 2007; Chong-Seng et al. 2014). With more frequent coral mass mortality events, post-mortality benthic composition may become a dominant state within tropical coral reefs, making it important to better understand reefs in this condition (Hughes et al. 2018; Emslie et al. 2024).

Coral reef benthic composition is inherently variable, with the identities and relative abundances of organisms varying with local environmental conditions (Sheppard 1980, Williams et al. 2013, Aston et al. 2019). Factors including depth, wind exposure, currents, and upwelling, are all known to affect the benthic composition of reefs (Sheppard 1980; Huston 1985; Gove et al. 2013; Williams et al. 2013; Gove et al. 2015; Williams et al. 2018; Aston et al. 2019; Ortiz et al. 2021). Mass coral bleaching and mortality can lead to further variability in benthic composition through spatial variation in the loss and recovery dynamics of coral cover (Golbuu et al. 2007; Green et al. 2008; Chong-Seng et al. 2014; Fox et al. 2019; Koester et al. 2020).

However, on most coral reefs, it is challenging to disentangle these sources of variability from the effects of local anthropogenic disturbances, such as coastal development (e.g. dredging), pollution, and overfishing, which also re-shape coral community composition (Anthony et al. 2015; Hoegh-Guldberg et al. 2018; Darling et al. 2019). Recognizing how natural environmental variability shapes post-mortality benthic composition is important for understanding the processes which shape reefs on a local scale.

Depth is an important source of natural variability in benthic composition, with light levels and the impact of waves decreasing with increasing depth. Greater depths can favour fleshy macroalgae whose photosynthetic pigments increase their ability to harvest light, as well as slow-growing massive and encrusting coral species, which can tolerate low light levels better than fast-growing branching and plating corals (Huston 1985; Darling et al. 2012; Williams et al. 2013). Similarly, wind exposure is a source of natural variation in benthic composition, as wind-induced wave action can increase turbidity and consequently reduce light penetration (Acevedo et al. 1989; Larcombe and Woolfe 1999). Moderate wind exposure can lead to coral communities dominated by slow-growing massive corals (Huston 1985; Darling et al. 2012; Maucieri and Baum 2021; Feliciano et al. 2023). Wave action itself can also favour massive or encrusting corals over branching and plating corals, and crustose coralline algae (CCA) over turf algae and fleshy macroalgae, because of their greater ability to withstand wave force (Madin 2005; Williams et al. 2013; Aston et al. 2019).

Depth and wave action can also lead to spatial variation in benthic composition following mass coral bleaching and mortality. Slow-growing massive and encrusting corals that dominate deeper sites have lower bleaching-driven mortality rates than the highly sensitive branching and plating corals that dominate the shallows (Bianchi et al. 2006; Darling et al. 2012; Gouezo et al. 2019). Wave action may also affect the recovery of corals post-mortality as recruitment may be higher at wave exposed sites where waves can reduce sediment and loosen rubble cover, making stable hard substrates, including CCA, more available for coral recruits (Tamelander 2002; Arthur et al. 2005; Chong-Seng et al. 2014; Yadav et al. 2016). Exposed conditions might also benefit recovering and newly recruited coral by reducing competition with turf and macroalgae, both of which are vulnerable to damage from waves (Loch et al. 2004; Fabricius 2005; Chong-Seng et al. 2014; Koester et al. 2020).

Here, we assessed factors contributing to local variation in benthic composition on Kiritimati (Republic of Kiribati), an isolated atoll in the central equatorial Pacific Ocean, following a coral mass mortality event in 2015-2016. Between 2014 and 2017, strong El Niño conditions led to one of the most severe global coral bleaching events ever recorded (Eakin et al. 2019). On Kiritimati, the El Niño caused 10 months of heat stress (June 2015 to April 2016), triggering mass coral bleaching and the loss of over 89% of hard coral cover (Baum et al. 2023) and all soft coral cover (Maucieri and Baum 2021) at sites along the 10-12 m isobath. Using benthic imagery, we characterized the benthic composition of shallow (7-10 m) and deep (18-22 m) reefs on the exposed and sheltered sides of the atoll three years post-mortality, and investigated how coral cover, algal cover, and abundance of juvenile coral (*i.*e., corals that recruited during or after the El Niño) vary spatially. We hypothesized that the effects of depth and wind exposure on benthic communities before, during, and after coral mass mortality would result in variation in post-mortality benthic composition. Specifically, we hypothesized that coral cover would be higher at deep and exposed locations, while juvenile coral colony abundance and CCA cover would be higher at shallow and exposed locations, and turf algae and fleshy macroalgae cover would be higher at deep and sheltered locations. We also hypothesized there would be an interaction between depth and wind exposure, since the strength of wave action decreases with depth, which would lead to weaker effects of wind exposure on benthic composition at deep exposed sites.

## Methods

### Study site and design

Data for this study were collected from the forereefs of Kiritimati, Republic of the Kiribati (01°52′N 157°24′W) in September 2019, three years after the end of the prolonged marine heatwave on the atoll that was induced by the 2015-2016 El Niño. This study examined 16 survey locations (spaced approximately 5 km apart) along the largely uninhabited southern coast of the atoll (**Figure 1**). Trade winds influence the eastern side of the island, such that half the locations (n = 8) were on the exposed (eastern) side of the island, while the other half (n = 8) were on the sheltered (western) side, as defined by the dominant southeasterly wind direction (Bosserelle et al. 2015; Claar et al. 2019). At the time, Kiritimati was home to 6500 residents, most of whom live in the island’s northwest region (Kiribati National Statistics Office 2016; Watson et al. 2016). This left the southern region of the island relatively undisturbed by human activity (Watson et al. 2016). For this study we did not directly quantify human disturbance level at our locations. However, previous studies have classified similar locations on Kiritimati as very low disturbance (Watson et al. 2016; Magel et al. 2020; Baum et al. 2023).

**Fig. 1.**
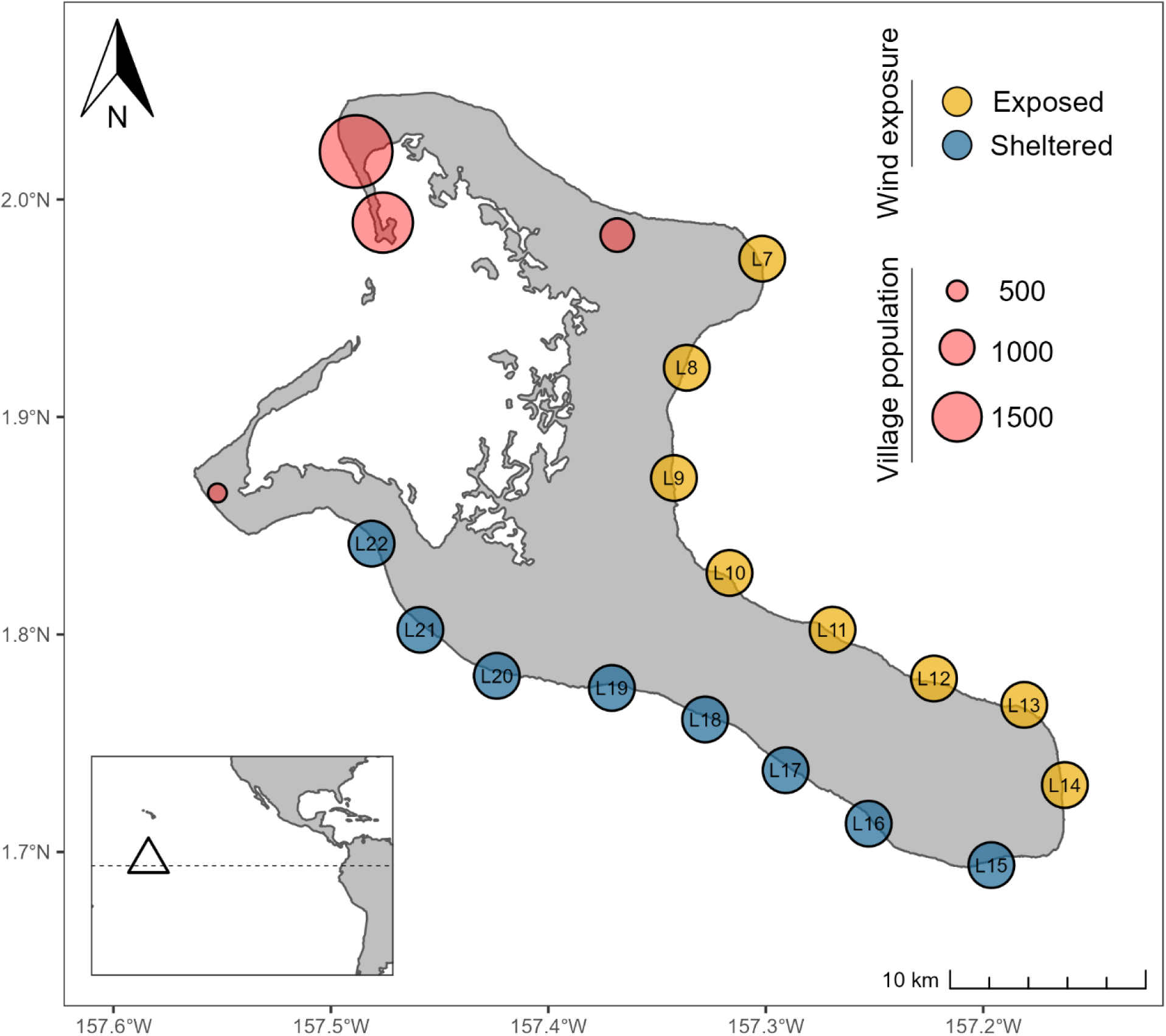
Map of Kiritimati Island, Kiribati with locations surveyed for this study. Exposed locations (L7-L14) are shown in yellow and sheltered locations (L15-L22) are shown in blue. Villages are shown by red circles with size proportional to human population size. The triangle in the inset indicates Kiritimati’s location in the central equatorial Pacific Ocean.

At each location, two sites were surveyed via SCUBA, one shallow site at 7-10 m and one deep site at 18-22 m (**Figure S1**). For each site, three 50 m transects were laid along the appropriate isobath, separated from one another by 5-10 m. Photos of the benthos were taken along these transects at intervals of approximately 1 m to create photo quadrats of 1 m^2^. A ruler was included in each photo for scale.

### Image analysis

For each of the 32 sites surveyed (16 locations with two sites at each), we randomly selected 16 photos from two separate transects to analyze. Only photos that were of useable quality were included in the random selection process. We cropped each of the selected photos to 90 x 60 cm using Fiji (Version 2.1.0), an open-source image processing package built on ImageJ software. For one site (L9 Deep) only eight photos were selected, as photos taken on two of the three transects were unusable. In total, 504 photos were randomly selected for analysis.

To quantify benthic composition, which included abiotic substrates as well as sessile organisms from the benthic community, we uploaded each cropped photo to CoralNet, an open-source image analysis software designed for annotating reef photo quadrats and identified the substrate under 54 randomly generated points (Beijbom et al. 2015). Each point was identified as one of 41 categories, which we further merged into nine substrate groupings for analysis: hard coral cover, soft coral cover, crustose coralline algae cover, *Peyssonelia* algae cover, fleshy macroalgae cover, turf algae cover, cyanobacteria cover, other biotic cover, and abiotic cover (**Table S1)**. Hard coral (order Scleractinia) were identified to the genus and species level whenever possible, and included 20 genera of reef-building corals and one family of free-living corals. Soft coral (order Alcyonacea) could only be identified to the genus level and included three genera. Fleshy macroalgae included Halimeda and *Caulerpa*, two genera of green (Chlorophyte) algae. Crustose coralline algae and cyanobacteria were not classified any further, while abiotic cover included sand, rubble, and dead coral skeleton.

For juvenile coral colony abundance, we defined juveniles as coral colonies who, based on size, health, and morphology, likely recruited during or after the 2015-2016 El Niño. We focused on two genera of corals that we observed most frequently and could classify as juveniles reliably*: Acropora* and *Pocillopora*. The El Niño led to bleaching and death for most *Acropora* and *Pocillopora* colonies on Kiritimati (Baum et al. 2023), so we expected the majority of colonies seen in 2019 photo quadrats to fit our definition of juvenile. To help exclude surviving adults from our count data, we did not include colonies with visual signs of regrowth or bleaching in counts. We also measured the diameter of each colony, so we could identify surviving adults based on size. As growth rates recorded for *Acropora* and *Pocillopora* vary widely between studies, we could not use growth rates to set a size limit for juveniles. Instead, we used the size distribution of colonies measured in this study to identify outliers in colony size which could represent surviving adults. One *Pocillopora* colony was found to be an outlier, as it had a diameter of 27 cm, while all other measured colonies (n = 263) had diameters less than 23 cm in diameter, and therefore was removed from further analysis (**Figure S2**). Due to our assessment of colony health and size to identify juvenile corals, we only counted colonies where at least 90% of their surface was visible within the photo quadrat.

### Statistical analysis

We performed all statistical analyses in RStudio with R version 4.3.1 (R Core Team 2023).

First, to examine the effect of depth and wind exposure on hard coral, CCA, fleshy macroalgae, and turf algae cover, we fitted generalized linear mixed-effects models (GLMMs) for mean proportion of cover for each substrate by transect. We did not model these effects for soft coral cover, because soft corals were rare overall and absent from most sites. For hard coral, CCA, and fleshy macroalgae cover we used a zero-inflated beta probability distribution (logit link function), which is appropriate for continuous proportion data containing zeroes, with the “glmmTMB” package (Brooks et al. 2017). For turf algae cover we used a beta probability distribution (logit link function) without zero-inflation since turf algae was rarely absent in our data. We modeled depth and wind exposure as categorical fixed effects, with shallow sites and sheltered locations as our baseline, and treated depth nested in location as a random intercept. Transect-level data allowed each location-depth combination to have multiple points for estimating a random intercept. We included models with and without an interaction term between depth and wind exposure. We compared models using Akaike Information Criterion for small sample sizes (AICc) in the “bbmle” package and reported all models within 6 AICc units of the top model (Bolker 2020). We visually assessed model assumptions using simulated residual plots from the “DHARMa” package (Hartig 2020). Additionally, we graphed coral cover by taxon, to examine the effects of depth and wind exposure on the presence and abundance of different coral species, in particular species with different life history strategies, as defined by Darling et al. (2012).

Next, to examine the effect of depth and wind exposure on juvenile *Acropora* and *Pocillopora* colony abundances, we fitted GLMMs for total colony count per transect for each genus. Since our count data were overdispersed, we used a negative binomial probability distribution (log link function) from the “glmmTMB” package (Brooks et al. 2017). We used the model structure as described for substrate cover and compared and validated models as above.

Finally, to examine the effect of depth and wind exposure on overall benthic composition, we performed multivariate analysis and ordination on cover by substrate. We calculated Bray-Curtis distances on a site by substrate grouping matrix and visualized these differences using Principal Coordinates Analysis (PCoA) from the “ape” package (Paradis and Schliep 2019). We tested for the effects of depth, wind exposure, and their interaction using permutational multivariate analysis of variance (PERMANOVA) with 999 permutations from the “vegan” package (Oksanen et al. 2020).

## Results

### Coral cover and community composition

Hard coral proportional cover was highest at deep sheltered sites (0.148 ± 0.014 (SE)) and lowest at shallow exposed sites (0.020 ± 0.003 (SE)), with an average proportion of 0.069 ± 0.005 (SE) across all sites and locations. (**Figure 2**). Depth and wind exposure were both significant predictors of hard coral cover (**Table 1, Table S2**), with deep and sheltered transects having significantly higher mean hard coral cover than shallow and exposed transects, respectively (**Table S2**, **Figure 3A**). The top model did not include an interaction between depth and wind, and when included in the second-best model, it was not statistically significant (**Table S2**). At all but one location, L16 (**Figure 1**), mean proportion of hard coral cover was higher at the deep site than the shallow site (**Figure 3B, Table S3**).

**Fig. 2.**
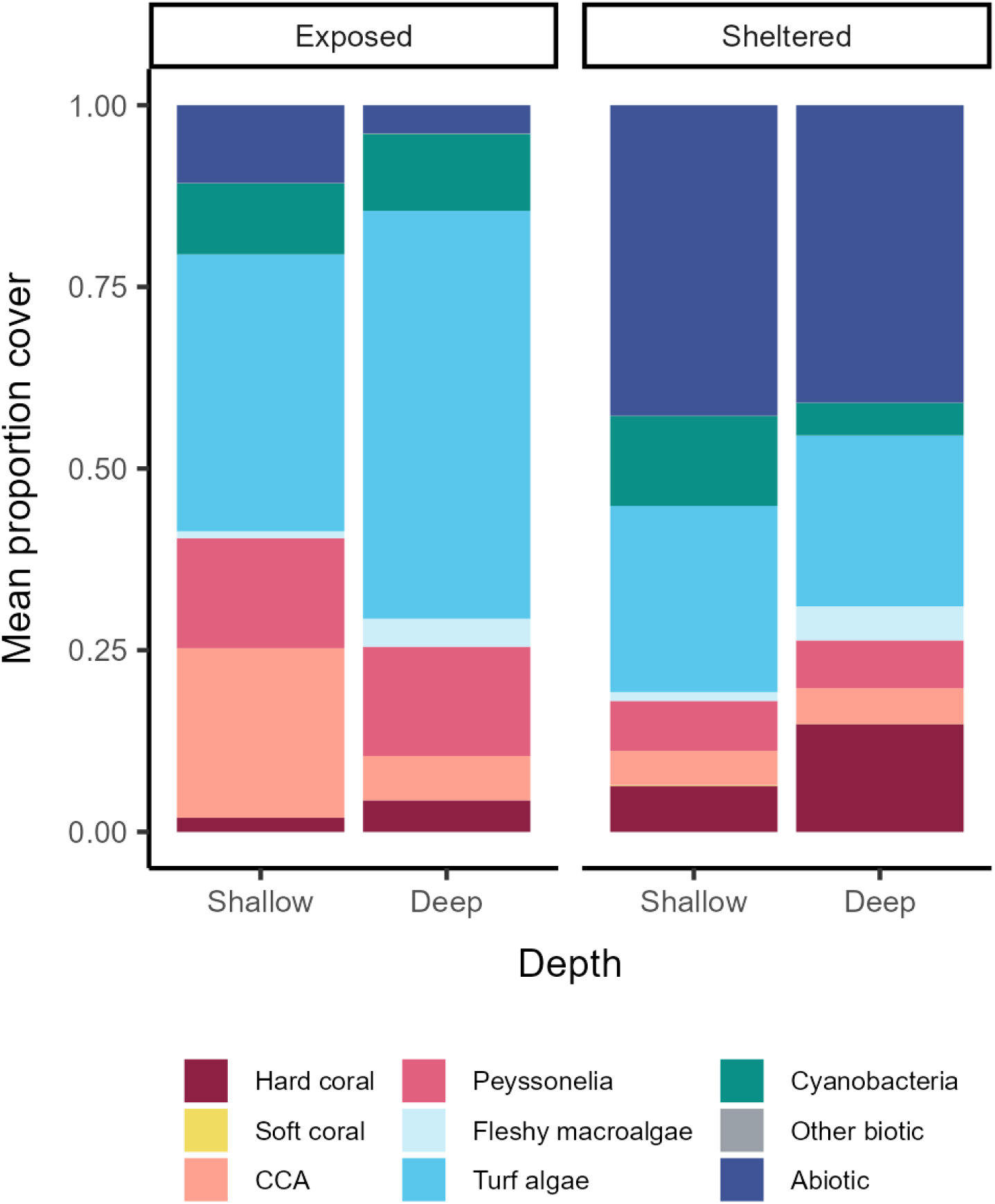
Barplots of mean proportion benthic cover by depth (shallow or deep) and wind exposure (exposed or sheltered). Benthic cover was classified into nine substrates: hard coral, soft coral, crustose coralline algae (CCA), encrusting macroalgae from the genus *Peyssonelia* (Peyssonelia), fleshy macroalgae, turf algae, cyanobacteria, other biotic substrates, and abiotic substrates. See Figure S3 for mean proportion cover by site.

**Figure 3.**
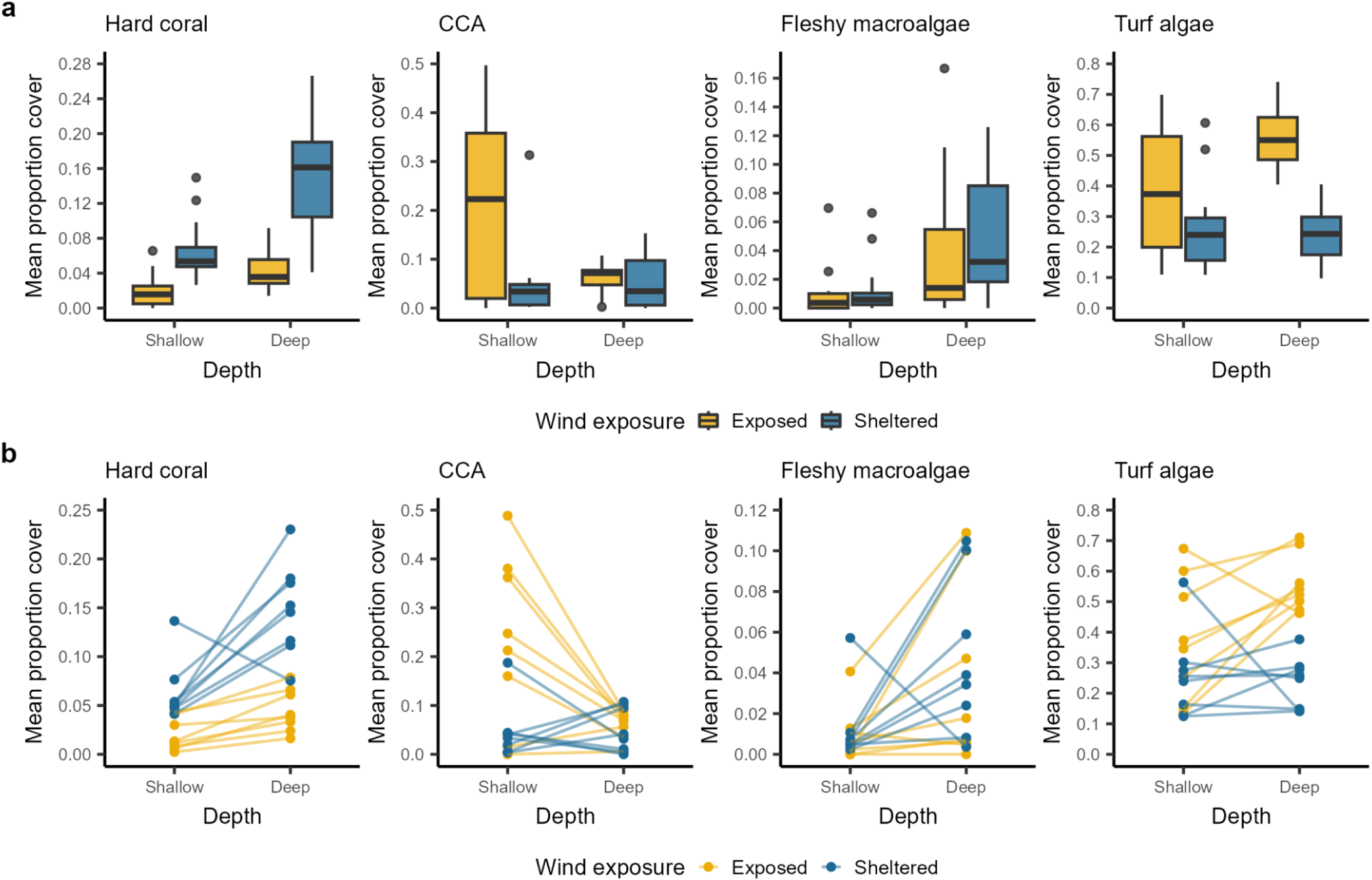
(a) Boxplot of mean proportion cover by substrate for transects from deep and shallow sites at exposed and sheltered locations. (b) Interaction plot of mean proportion cover by substrate from deep and shallow sites at exposed (yellow) and sheltered (blue) locations. To improve visualization, two large outliers were removed from the boxplots. Note that y-axes differ between panels. See Tables S3-S7.

**Table 1.** Model summaries for generalized linear mixed effects which examined the effects of depth and wind exposure on the mean proportion of hard coral, CCA, fleshy macroalgae, and turf algae cover, and the total number of *Acropora* and *Pocillopora* colonies. Only models within 6 AIC units of the top model are reported. Model terms which are significantly different from zero are bolded.

| Response | Model | $\Delta\text{AICc}$ | df | $w_i$ |
| --- | --- | --- | --- | --- |
| Hard coral cover | **Depth + Wind** | 0.0 | 7 | 0.770 |
|  | **Depth + Wind** + Interaction | 2.4 | 8 | 0.230 |
| CCA cover | Depth + **Wind** + **Interaction** | 0.0 | 8 | 0.833 |
|  | Depth + **Wind** | 4.2 | 7 | 0.100 |
|  | **Wind** | 5.2 | 6 | 0.061 |
| Fleshy macroalgae cover | **Depth** | 0.0 | 6 | 0.700 |
|  | **Depth** + Wind | 2.3 | 7 | 0.220 |
|  | **Depth** + Wind + Interaction | 4.7 | 8 | 0.067 |
| Turf algae cover | Depth + Wind + **Interaction** | 0.0 | 7 | 0.523 |
|  | Wind | 1.5 | 5 | 0.243 |
|  | Depth + **Wind** | 1.6 | 6 | 0.233 |
| *Acropora* count | **Depth** | 0.0 | 5 | 0.686 |
|  | **Depth** + Wind | 2.1 | 6 | 0.244 |
|  | **Depth** + Wind + Interaction | 4.6 | 7 | 0.069 |
| *Pocillopora* count | Depth | 0.0 | 5 | 0.596 |
|  | Depth + Wind | 2.4 | 6 | 0.180 |
|  | Wind | 2.9 | 5 | 0.140 |
|  | Depth + Wind + Interaction | 3.9 | 7 | 0.083 |

Stress-tolerant hard coral taxa made up the majority of coral cover (**Figure 4**), with genera such as *Porites, Pavona, Dipsastraea,* and *Platygyra* representing more than 60% of the total coral cover. In contrast, competitive hard corals (*Pocillopora, Acropora, Turbinaria,* and *Montipora*) made up less than 10% (**Table S4**). At sheltered locations, *Porites* had the highest average proportion of cover, while at exposed locations *Pavona durdeni* was more prevalent (**Figure 4**). Although stress-tolerant corals had the greatest mean proportion cover at all depth and exposure levels, competitive corals were relatively important at exposed locations. *Acropora* was the second most abundant coral taxon at shallow exposed sites, while *Pocillopora* was the third-most abundant taxon at deep exposed sites. The proportion of soft coral cover was less than 0.001 ± 0.000 (SE) when averaged across all sites and locations. Soft corals were only found at sheltered locations, making up less than 2% of all coral cover at shallow sites, and less than 0.5% at deep sites (**Figure 2**).

**Fig. 4.**
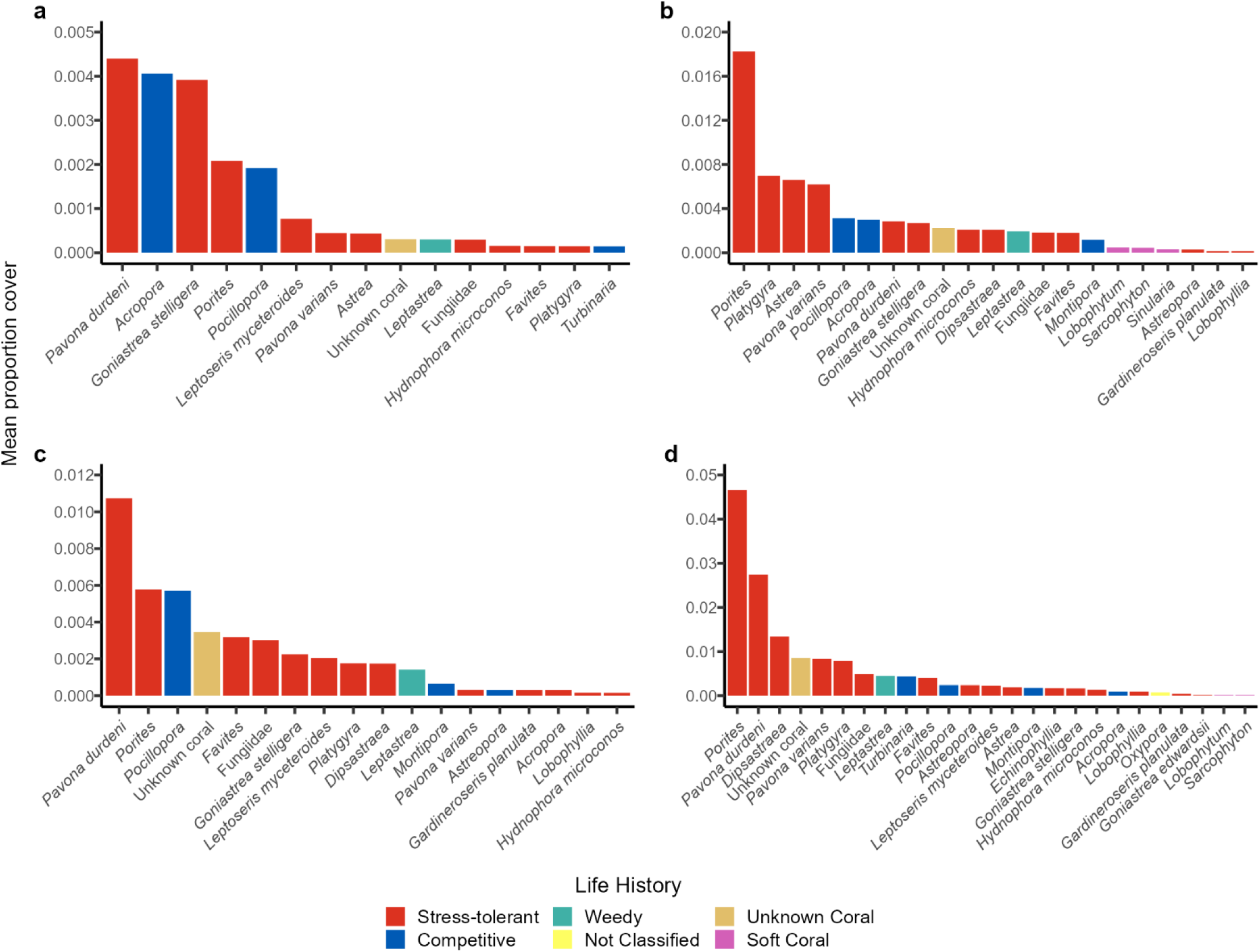
Barplots of mean proportion coral cover by taxon for a) shallow exposed sites b) shallow sheltered sites c) deep exposed sites and d) deep sheltered sites. Note that y-axes differ between panels. Life history classifications (following Darling et al. 2012) are included for each hard coral taxon. See Figure S4 for mean proportion cover across all sites.

### Juvenile hard coral colony abundance

In total, we observed 105 juvenile *Acropora* colonies and 158 juvenile *Pocillopora* colonies across all sites. Mean *Acropora* colony abundance was highest at exposed and shallow sites (0.367 ± 0.072 (SE) per site) and lowest at sheltered and deep sites (0.086 ± 0.029 (SE) per site). Mean *Pocillopora* colony abundance was highest at exposed and deep sites (0.492 ± 0.086 (SE) per site) and lowest at exposed and shallow sites (0.156 ± 0.039 (SE) per site).

Depth was the single best predictor of juvenile *Acropora* colony abundance (**Table 1, Table S5**). Deep transects had significantly lower *Acropora* counts than shallow transects, with only one location, L12 (**Figure 1**), having equal mean *Acropora* counts for the shallow and deep sites (**Table S6, Figure 5A & 5B)**. All other locations had lower mean *Acropora* counts at deep sites relative to shallow sites at the same location.

**Fig. 5.**
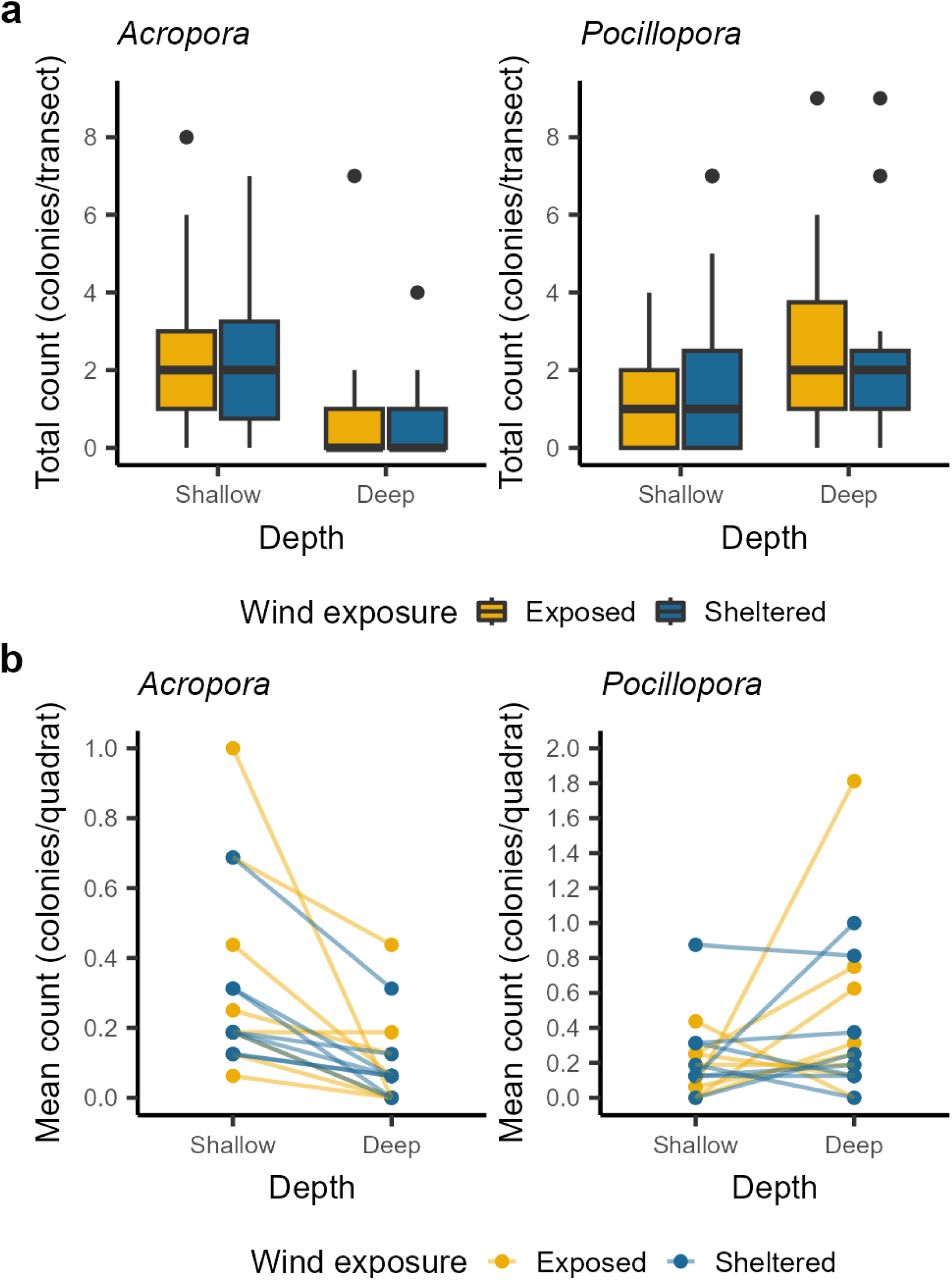
(a) Boxplot of total *Acropora* and *Pocillopora* colony counts per transect from deep and shallow sites at exposed and sheltered locations. (b) Interaction plot of mean *Acropora* and *Pocillopora* colony counts per quadrat from deep and shallow sites at exposed and sheltered locations. Exposed locations are shown in yellow, while sheltered locations are shown in blue. To improve visualization, three large outliers were excluded from the boxplots. See Table S9 for a list of mean count by site.

Depth was also the only fixed effect included in the best model for juvenile *Pocillopora* colony abundance, but the effect was not significant (**Table 1**, **Table S4**). Depth was also not significant in any of the other selected models. At the location level, differences in mean colony counts between shallow and deep sites were inconsistent. Nine locations had higher mean counts at deep sites, five locations had higher mean counts at shallow sites, and two locations had no difference between shallow and deep sites (**Figure 5B**).

There was no consistent effect of wind exposure on *Acropora* or *Pocillopora* colony counts (**Figure 5A & 5B**). Wind exposure was a poor predictor of *Acropora* and *Pocillopora* abundance, since its effects were not significant in models where it was included as a predictor (**Table S4**). Interactions between wind exposure and depth were also not significant in models for either coral genus.

### Crustose coralline, macroalgae, and turf algae cover

On average, crustose coralline algae (CCA) represented 0.098 ± 0.007 (SE) of benthic cover, although, at shallow and exposed sites, where CCA cover was highest, this proportion more than doubled (**Figure 2, Table S7**). Wind exposure was a significant predictor of CCA cover in the top model, which predicted higher CCA cover for exposed locations (**Table 1**, **Table S2**). Depth was also included in the top model, however, it was not a significant predictor of CCA cover (**Table 1**, **Table S2**). In the other two selected models, wind was a significant predictor while depth was either not significant or not included in the model. While the median CCA cover at shallow exposed locations was greater than median cover at shallow sheltered locations, there was very little difference in CCA cover at deep exposed locations compared to deep sheltered locations (**Figure 3A**). This reflects the significant interaction between depth and wind reported by the top model (**Table 1, Table S2)**. Except for L8 and L13 (**Figure 1**), CCA cover was higher for shallow sites than deep sites at exposed locations, while at sheltered locations, there was no clear effect of depth (**Figure 3B, Table S8**).

Fleshy macroalgae represented only 0.027 ± 0.003 (SE) of benthic cover when averaged across all sites and locations. At both exposed and sheltered locations, macroalgae cover was higher at deep than at sheltered sites (**Table S7, Figure 2**). Depth was a significant predictor in the top model, with a positive effect on fleshy macroalgae cover (**Table 1**, **Table S2**). While macroalgae cover ranged widely for both deep and shallow sites, all but two locations, L9 and L16 (**Figure 1**), had greater macroalgal cover at their deep sites relative to their shallow sites (**Figure 3A & 3B, Table S9**). Neither the effect of wind nor the interaction between wind and depth were significant for macroalgal cover.

Overall, turf algae were the most abundant substrate, with an average proportion of 0.355 ± 0.010 (SE). Turf algae cover was even greater at exposed locations, while at sheltered locations, abiotic cover replaced turf algae as the most abundant substrate (**Table S7**, **Figure 2**). For the top model selected for turf algae cover, the effect of wind exposure was not significant but the interaction between wind and depth was (**Table 1, Table S2**). In the other selected models, however, which did not include an interaction term, wind exposure was a significant predictor. Turf cover for deep exposed locations was consistently greater than turf cover for deep sheltered locations (**Figure 3A**). While there was a greater range in turf cover at shallow sites, cover was higher for shallow exposed locations compared to shallow sheltered locations. Depth was not a significant predictor of turf cover. However, for all exposed locations except L8 (**Figure 1**), turf cover was higher for the deep site compared to the shallow site (**Figure 3B, Table S10**). There was no pattern in turf cover with depth for sheltered locations.

### Variation in benthic composition

Benthic composition, defined as both the biotic and abiotic components of benthic cover, differed significantly with wind exposure, with wind explaining 57 % of the variation in composition (**Table 2**, **Figure 6**). While depth explained 27% of variation, the effect was not significant, and neither was the interaction between depth and wind. Accordingly, there was very little overlap in ordination space between exposed and sheltered locations (**Figure 6**). There was, however, considerable overlap between deep and shallow sites, especially for sheltered locations.

**Fig. 6.**
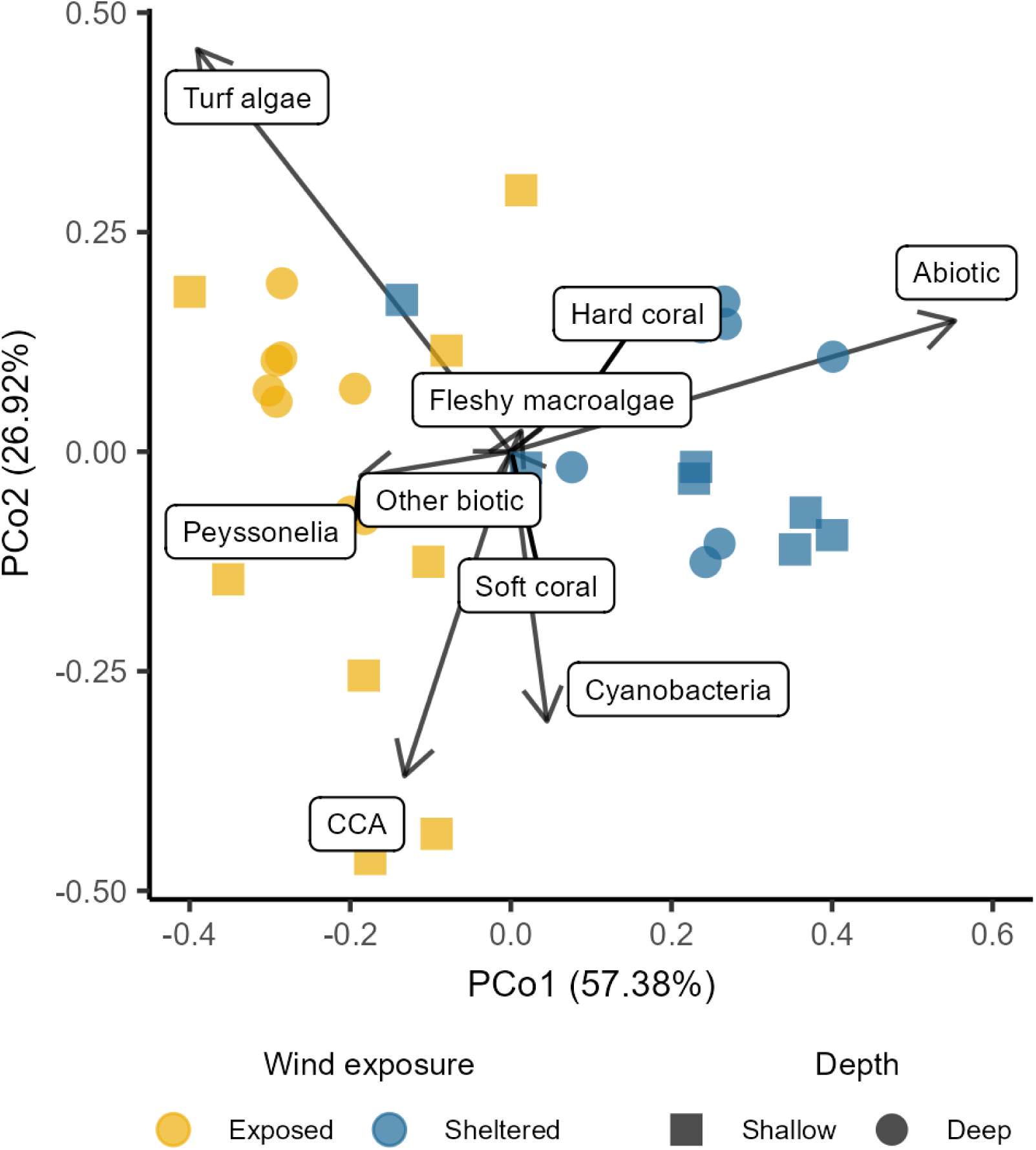
Biplot of principle coordinates analysis (PCoA) for benthic composition, quantified as Bray-Curtis dissimilarity. Shallow (square) and deep (circle) sites at exposed (yellow) or sheltered (blue) locations are shown relative to the first and second PCoA axes. Arrows represent the nine types of benthic cover used to calculate the dissimilarity matrix.

**Table 2.** Results of PERMANOVA test examining the effects of depth and wind exposure on benthic composition. Bolding indicates significance.

| Source | df | Sum of Squares | Pseudo-F | R<sup>2</sup> | *P* (perm) |
| --- | --- | --- | --- | --- | --- |
| Depth | 1 | 0.1612 | 2.7465 | 0.04712 | 0.051 |
| Wind | 1 | 1.4869 | 25.3389 | 0.43474 | **0.001** |
| Depth:Wind | 1 | 0.1291 | 2.2004 | 0.03775 | 0.105 |
|  |  |  | * *P* < 0.05 | ** *P* < 0.01 | *** *P* < 0.001 |

## Discussion

Three years after a severe coral mass mortality event, hard and soft coral cover were low on the low-disturbance reefs of Kiritimati, representing less than 10% of total benthic cover on average. Hard coral cover varied spatially, with higher cover at exposed locations and deep sites. Stress-tolerant hard corals made up the majority of coral cover, while juvenile colonies of competitive hard corals were less important, except at wind exposed sites where stress-tolerant corals were less abundant. Although the exact sites within this study were not surveyed before 2019, coral cover at nearby shallows sites was 55.1% prior to the 2015-2016 El Niño and had declined to 3.3% by 2017 (Baum et al. 2023). In 2019, coral cover at the shallow sites surveyed in the present study was 4%, suggesting that there was very little coral recovery between 2017 and 2019 (**Table S11**). While cover by non-coral substrates also varied spatially on Kiritimati’s low-disturbance post-mortality reefs, these patterns were consistent with increased substrate availability following coral mass mortality, and substrate-specific preferences in terms of abiotic conditions. We suggest that pre-mortality coral composition as well as post-mortality recruitment were responsible for these spatial patterns in benthic community composition.

### Coral cover and community composition

Differences in pre-mortality coral community composition best explained the post-mortality spatial distributions of coral that we observed. Pre-mass mortality, large competitive colonies, especially adult tabulate *Acropora*, dominated reefs studied on the exposed side of Kiritimati but were less abundant at sheltered locations, while stress-tolerant and massive coral species were reported on the exposed and sheltered sides of Kiritimati (Tietjen 2020; Baum et al. 2023). Slow-growing species with massive or encrusting morphologies are often stress-tolerant, meaning they survive bleaching events at relatively high rates (Baird and Marshall 2002; Darling et al. 2012). Fast-growing competitive species with branching and plating morphologies, however, survive at much lower rates (Baird and Marshall 2002; Darling et al. 2012; Depczynski et al. 2013; Pisapia et al. 2016; Head et al. 2019). Therefore, higher competitive coral cover at exposed locations would have meant a greater rate of coral mortality, and subsequently, lower post-mortality coral cover relative to sheltered locations.

The composition of coral communities we observed in 2019 also supported pre-mortality composition as the major driver of post-mortality cover. The majority of corals we recorded were large colonies of stress-tolerant mounding and encrusting species, since nearly all of the competitive corals on Kiritimati died during the coral mass mortality (Tietjen 2020; Baum et al. 2023). The branching and plating corals that we observed were largely *Acropora* and *Pocillopora* juveniles, which contributed less to coral cover than large, stress-tolerant survivors, because of their small size and low abundance. While competitive corals made up a larger proportion of coral cover at exposed and sheltered locations, counts of juvenile *Acropora* and *Pocillopora* colonies were similar, indicating that their contribution to total cover was only large relative to the very low numbers of surviving of adult corals at these sites.

Similarly, the differences in coral cover with depth that we reported were consistent with this relationship between pre-mortality coral composition and post-mortality coral cover. While pre-mortality research on Kiritimati has only documented coral composition at shallow sites, stress-tolerant corals are known to make up a greater proportion of the coral community at deeper depths (Huston 1985; Jackson 1991; Williams et al. 2013). Stress-tolerant coral cover was likely higher at deep sites than shallow sites pre-mortality, resulting in lower mortality and therefore higher coral cover at deep sites post-mass mortality.

Overall, the post-mortality coral cover on Kiritimati was largely driven by adult stress-tolerant corals that survived the coral mass mortality event. This is consistent with research on other isolated reef systems where local recruitment rates were low in the first few years of recovery. In the absence of new juvenile coral colonies, post-mortality coral cover is mostly the product of the survival and regrowth of adult coral colonies (Arthur et al. 2005; Gilmour et al. 2013; Koester et al. 2020). Under these circumstances, spatial variation in post-mortality benthic composition is largely shaped by variation in pre-mortality coral composition, which results from differences in environmental conditions (Koester et al. 2020; Kobluk et al. 2021). In contrast, when low disturbance reefs are experiencing high recruitment, spatial variation in recruitment determines benthic composition on post-mortality reefs (Holbrook et al. 2018; Koester et al. 2021; Tebbett et al. 2022).

Unlike with hard corals, post-mortality soft coral distributions reflected the sensitivity of juvenile colony recruitment and survival to local environmental conditions. Maucieri and Baum (2021) reported 100% mortality of soft coral on Kiritimati following the 2015-2016 coral mass mortality event, with annual surveys of the shallow forereefs revealing no soft corals until 2019. Therefore, the soft corals observed in our study likely represent newly established colonies. We found that soft coral was only present at sheltered locations, which is consistent with soft corals’ sensitivity to wave action (Cornish and DiDonato 2004; Spencer et al. 2005; Maucieri and Baum 2021). However, given that soft corals represented less than 1% of total coral cover, these patterns were not drivers of the spatial variation in post-mortality coral cover reported in this study.

### Variation in algae cover

Previous studies have described turf algae growth as opportunistic and have reported very high abundances of turf following coral mass mortality, which is consistent with our observations on Kiritimati (Vargas-Ángel et al. 2019; Koester et al. 2020). The spatial patterns we observed in turf algae cover likely represented the inverse of patterns in abiotic cover. Turf algae cover was higher at exposed sites, where abiotic cover was lower, and vice versa.

However, turf algae are capable of growing under low to moderate sediment cover, and can even increase sediment levels by trapping particulate matter (Fabricius and De’ath 2001). This suggests that turf algae cover was abundant at both sheltered and exposed locations, but at sheltered locations was covered by sediment and therefore recorded as abiotic cover.

Together, abiotic cover and turf algae cover made up the majority of the benthos regardless of wind exposure or depth. However, at certain sites CCA and macroalgae cover was also relatively high. CCA was abundant at shallow exposed sites, where high wave exposure and high light intensity could have given CCA a competitive advantage over turf algae (Huston 1985; Bessell-Browne et al. 2017; Tâmega and Figueiredo 2019; Aston et al. 2019). Fleshy macroalgae cover, meanwhile, was highest at deep sites. The majority of macroalgae we documented was from the genus *Halimeda*, which are well adapted to low light conditions (Williams et al. 2013; Chong-Seng et al. 2014). This specialization could make it more able to compete against turf algae at greater depths (Williams et al. 2013).

Competition between algae and coral has long been suggested as a source of spatial variation in coral cover following coral mass mortality (e.g. Chong-Seng et al. 2014, Koester et al. 2020). Given that competition with algae most affects juvenile coral colonies, and we found that coral cover was mostly attributable to surviving adult colonies, spatial patterns in juvenile abundance could not have explained the variation in coral cover that we observed. In our study, patterns in coral cover and turf algae cover likely represent opportunistic growth by turf algae following loss of coral cover, as seen on other post-mortality reefs (Tebbett et al. 2022; Huntington et al. 2022; Khen et al. 2022).

### Limitations

Analyses in this study were limited by a lack of benthic composition data from the study sites before the coral mass mortality event. However, previous surveys of Kiritimati’s coral reefs by Walsh (2011) and Baum et al. (2023) documented pre-mortality benthic composition within 1 km of both exposed and sheltered locations used in this study (**Figure S5**). This means we were able to ground our findings of with spatially comparable data, which suggested that the variation in benthic composition with depth and wind exposure that we observed was best explained by spatial variation in coral mortality rates.

### Conclusions

Three years after a prolonged marine heatwave caused mass coral mortality on Kiritimati’s reefs, hard coral cover was very low (0.069 ± 0.005 (SE)) across sites with low local anthropogenic impacts. While hard coral was more abundant at deep sites and sheltered locations, all locations showed dramatic declines from pre-mass mortality coral cover reported from the same study region (Baum et al. 2023). Our results suggest that variation in adult coral mortality was the most important factor in post-mortality benthic composition.

While there are currently very few signs of different recovery trajectories between depths or wind exposure levels on Kiritimati, this could change in future. For example, low recruitment early in recovery can lead to algal dominance later in a reef’s recovery trajectory (Graham et al. 2015). Continued monitoring is required to determine whether turf and macroalgae will prevent a coral-dominated state as coral recruitment and competition for space increases in the future. Despite low juvenile coral abundances, it is encouraging that both *Acropora* and *Pocillopora* juveniles are present on the reef, given such high losses of adult *Acropora* and *Pocillopora* during the mass mortality event. If these juveniles survive to reproduce, the fast growth rates of their young may allow for rapid repopulation, as previously recorded on other reefs several ro many years post-mass mortality (Halford et al. 2004; Arthur et al. 2005; Gilmour et al. 2013). This could trigger another shift in benthic composition, as exposed locations and shallow sites, where competitive corals like *Acropora* and *Pocillopora*, previously dominated, may regain coral cover over time (Ortiz et al. 2021).

With time and low disturbance, Kiritimati’s reefs have the potential to recover regardless of depth and wind exposure, as other reef systems have (e.g. Golbuu et al. 2007, Pisapia et al. 2016, Gouezo et al. 2019, Emslie et al. 2024). However, given the increasing frequency of heat stress events, it is unclear whether there will be enough time for this recovery to occur. If Kiritimati experiences another mass mortality event before populations of competitive corals recover, this could permanently prevent a return to pre-mortality benthic composition. In light of this, our findings reaffirm the importance of addressing anthropogenic climate change and facilitating the recovery of coral reefs.

## Statements and Declarations

### Competing Interests

The authors have no competing interests to declare.

### Data Availability Statement

Data are available via the following URL: https://github.com/rghansen/Hansen_etal_2025_CoralReefs

## Acknowledgements

R.L.G.H. acknowledges funding through a University of Victoria JCURA award and an NSERC Graduate Scholarship – Master’s. D.G.M. was funded by the University of Victoria and NSERC Graduate Scholarships – Master’s and Doctoral. We gratefully acknowledge Megan Dethier, Rebecca Maher, Kindall Murie, and Olivia Graham for reviewing an early version of the manuscript.

## Author’s Contributions

A.H. designed field surveys and collected the data. J.K.B. and R.L.G.H. conceived the analytical ideas. R.L.G.H and D.G.M. processed and analyzed the data. R.L.G.H. wrote the first draft of the manuscript. All authors contributed to editing the manuscript and gave final approval for publication.

## Notes

### Competing Interest Statement

The authors have declared no competing interest.

