## Supplemental Information for "Spatial variation in benthic community composition on a minimally disturbed coral reef in the years following a prolonged marine heatwave"

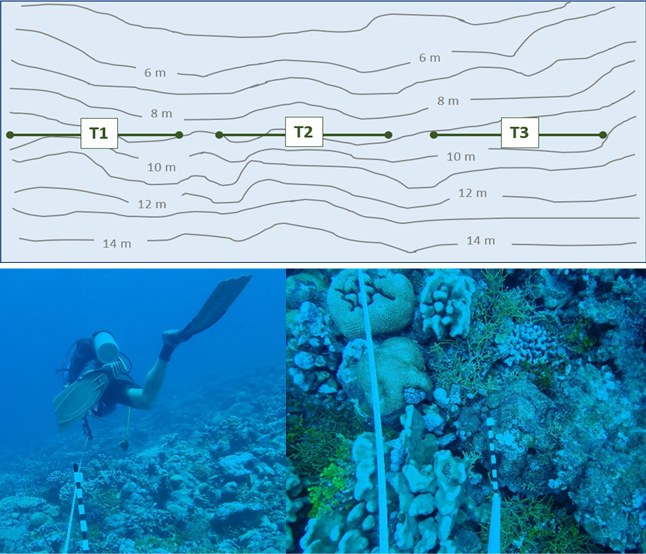

### Supplementary information

**Figure S1**. Transect-based surveys of the reef used to produce benthic photo quadrats. At each site, three transects were placed at along a single isobath, either between 7 and 10 m at shallow sites, or between 18 and 22 m at deep sites. Photos of the benthos were taken along each transect at intervals of approximately 1 m. Each photo included a ruler for scale.

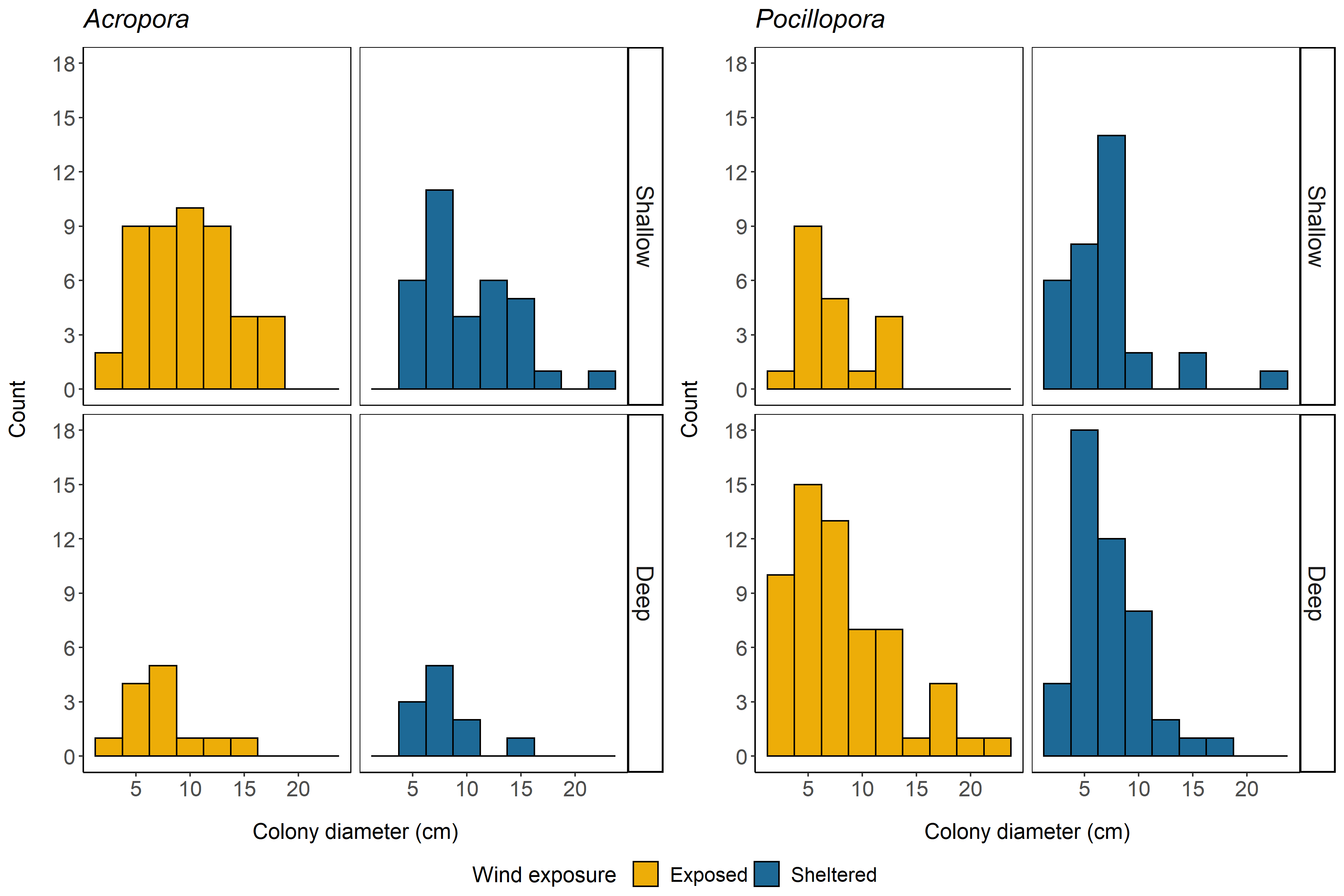

**Figure S2**. Histograms of *Acropora* and *Pocillopora* colony diameters for deep and shallow sites at exposed and sheltered locations.

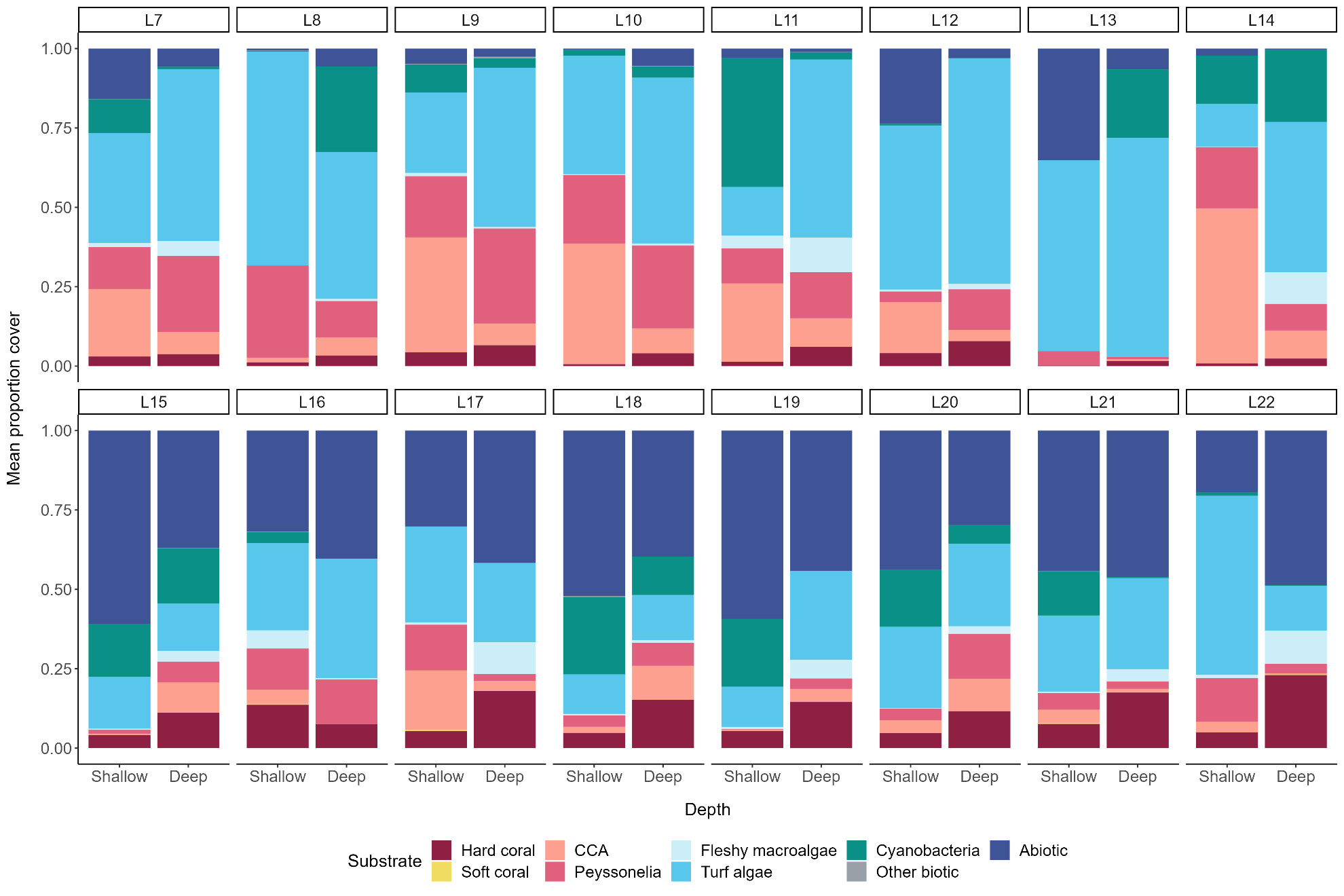

**Figure S3**. Barplots of mean proportion benthic cover for shallow and deep sites at exposed (first row) and sheltered (second row) locations. Benthic cover was classified into nine substrates: hard coral, soft coral, crustose coralline algae (CCA), encrusting macroalgae from the genus *Peyssonelia* (Peyssonelia), fleshy macroalgae, turf algae, cyanobacteria, other biotic substrates, and abiotic substrates.

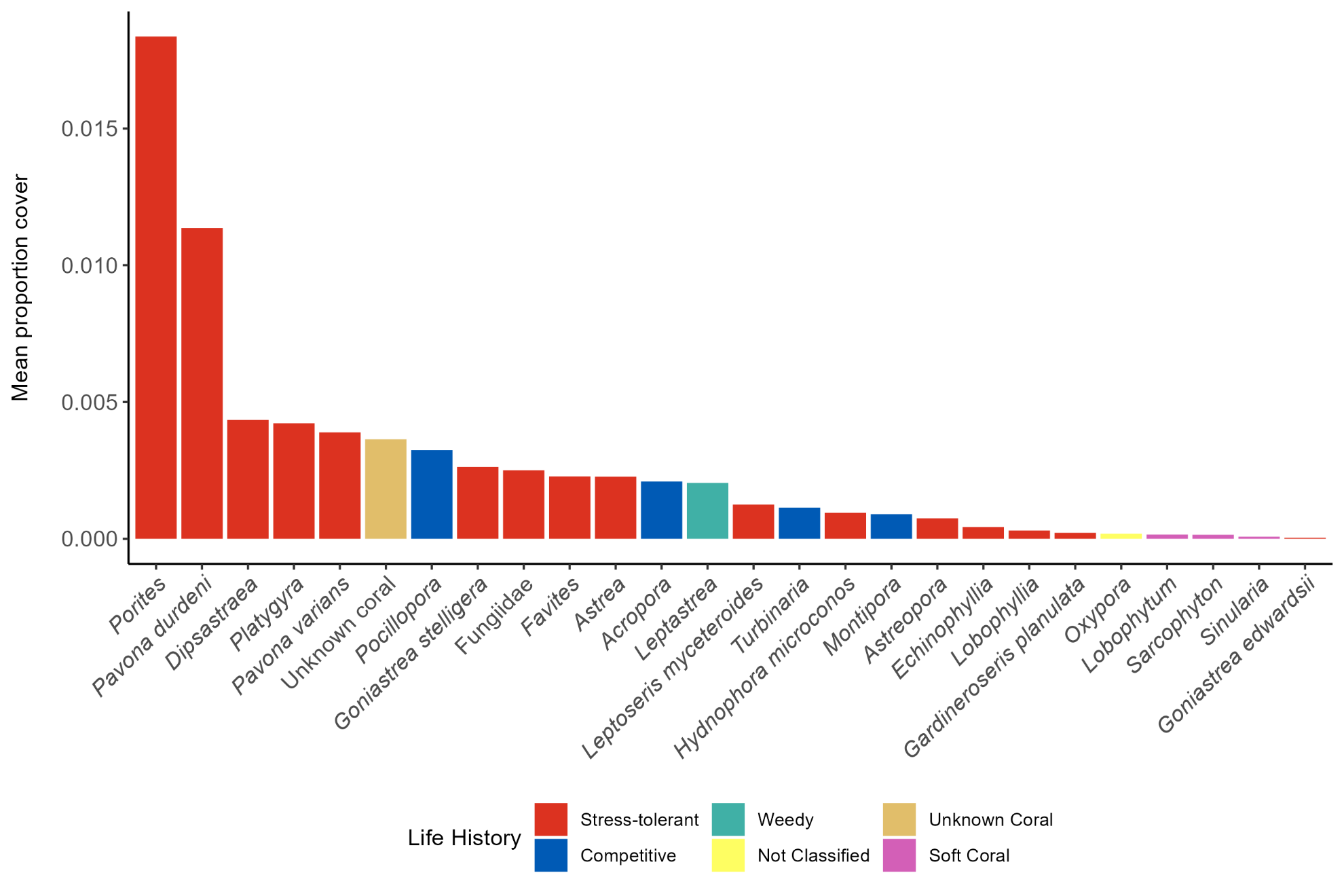

**Figure S4**. Barplot of mean proportion coral cover by taxon across all sites. Life history classifications (following Darling et al. 2012) are included for each hard coral taxon.

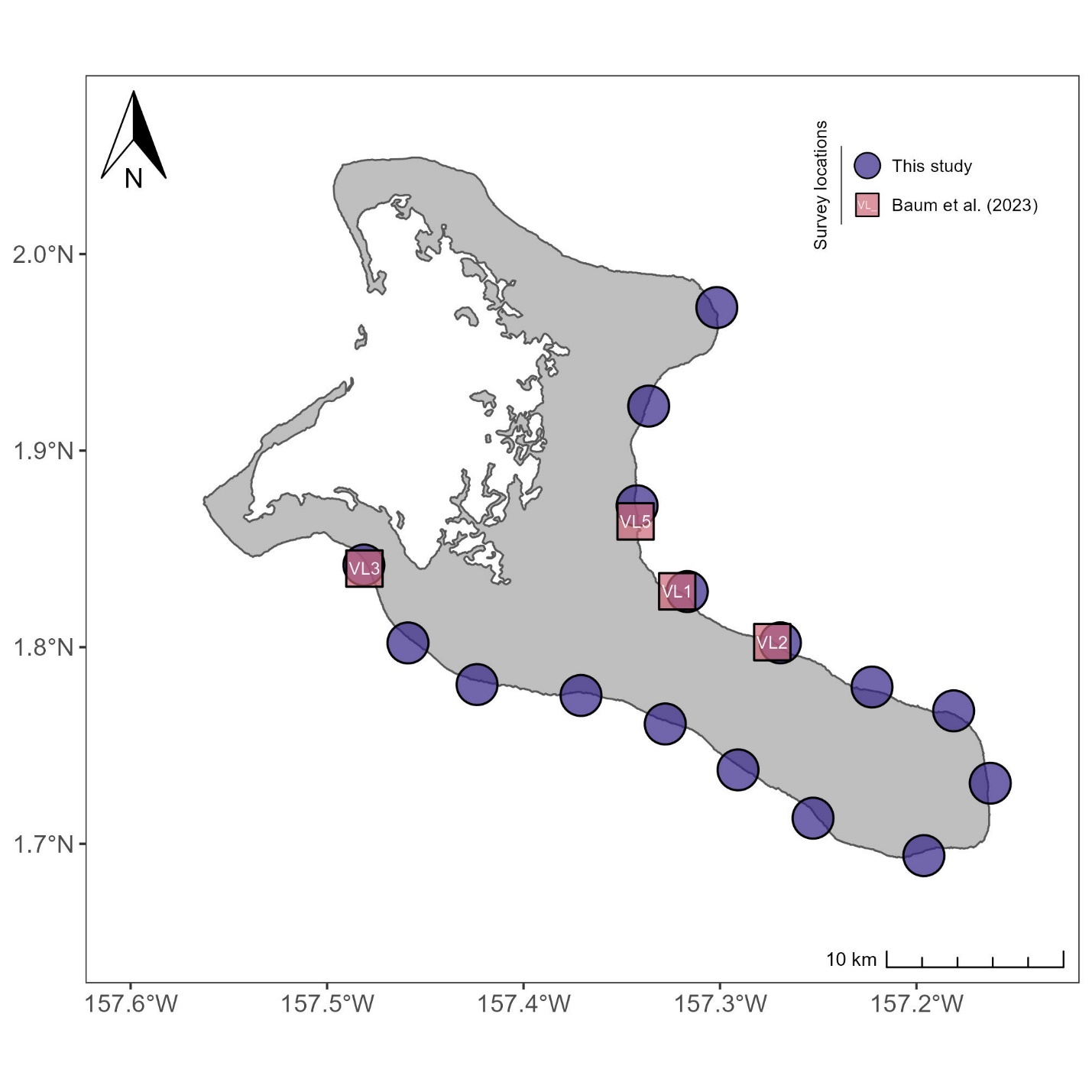

**Figure S5**. Locations surveyed by Baum *et al.* 2023 relative to those surveyed in this study. Baum *et al.* 2023 collected reef benthic composition data before and shortly after the 2015-2016 caused a coral mass mortality event on Kiritimati.

| **Substrate Grouping** | **Category** | **Definition** |
| --- | --- | --- |
| Abiotic | Rubble | Unattached hard substrate |
|  | Consolidated rock | Exposed dead coral skeleton |
| **Table S1**. Categories included in each substrate grouping used to classify benthic cover. | Sand | Sand |
|  | Sediment | Sand covering hard substrate |
| CCA | Crustose coralline algae (CCA) | Crustose coralline algae (CCA) |
| Cyanobacteria | Cyanobacteria film | Cyanobacteria |
| Hard coral | *Acropora* | All branching and tabulate forms of *Acropora* seen on Kiritimati |
|  | *Astrea* | *Astrea annuligera, A. curta* |
|  | *Astreopora* | *Astreopora cucullata, A. listeri, A. myriophthalma, A. scabra, A. suggesta, A. expansa, A. gracilis, A. ocellata*. |
|  | *Dipsastraea* | *Dipsastraea favus, D. laxa, D. matthaii, D. pallida, D. rotumana, D. speciosa* |
|  | *Echinophyllia* | *Echinophyllia aspera* |
|  | *Favites* | *Favites chinensis, F. abdita, F. complanata, F. flexuosa, F. halicora, F. pentagona* |
|  | Fungiidae | *Sandalolitha dentata, S. robusta, Herpolitha limax, H. weberi, Ctenactis echinata, Cycloseris cyclolites, Danafungia scruposa, D. horrida, Fungia fungites, Lithophyllon repanda, L. concinna, Lobactis scutaria, Pleuractis granulosa, P. paumotensis* |
|  | *Gardineroseris planulata* | *Gardineroseris planulata* |
|  | *Goniastraea edwardsii* | *Goniastraea edwardsii* |
|  | *Goniastraea stelligera* | *Goniastrea stelligera* |
|  | *Hydnophora microconos* | *Hydnophora microconos* |
|  | *Leptatstrea* | *Leptastrea* spp. |
|  | *Leptoseris myceteroides* | *Leptoseris myceteroides* |
|  | *Lobophyllia* | *Lobophyllia corymbosa, L. hataii, L. hemprichii, L. robusta* |
|  | *Montipora* | All foliose and encrusting forms of *Montipora* seen on Kiritimati Island |
|  | *Oxypora* | *Oxypora* spp. |
|  | *Pavona duerdeni* | *Pavona duerdeni* |
|  | *Pavona varians* | *Pavona varians*, but also may include *P. chiriquiensis* and *P. venosa* |
|  | *Platygyra* | *Platygyra contorta, P. daedalea, P. lamellina, P. pini, P. ryukyuensis, P. sinensis* |
|  | *Pocillopora* | All branching forms of *Pocillopora* seen on Kiritimati Island |
|  | *Porites* | *Porites australiensis, P. lichen, P. lobata, P. lutea, P. lukoensis, P. solida, P. superfusa, P. vaughani* |
|  | *Turbinaria* | All foliose forms of *Turbinaria* seen on Kiritimati Island |
|  | Unknown Coral | Can be identified as hard coral but there is not enough detail to be able to identify it to any lower taxon |
| Fleshy macroalgae | *Caulerpa* | *Caulerpa* spp. |
|  | *Halimeda* | *Halimeda* spp. |
|  | Macroalgae | Fleshy macroalgae that cannot be identified to genus |
| *Peyssonnelia* | *Peyssonnelia* | *Peyssonnelia* spp. |
| Other biotic | *Palythoa sp* | *Palythoa* spp. |
|  | Sea cucumber | Sea cucumber |
|  | Sea urchin | Sea urchin |
|  | Sponge | Sponge |
| Soft coral | *Lobophytum* | *Lobophytum* spp. |
|  | *Sarcophyton* | *Sarcophyton* spp. |
|  | *Sinularia* | *Sinularia* spp. |
| Turf algae | Turf algae | Turf algae |

| Response | | Model | | AICc | | ΔAICc | | df | *w_i_* | | Predictor | | | Estimate | | | SE |
| --- | --- | --- | --- | --- | --- | --- | --- | --- | --- | --- | --- | --- | --- | --- | --- | --- | --- |
| Hard coral cover | | Depth + Wind | | - 231.6 | | 0.0 | | 7 | 0.770 | | Intercept | | | **- 2.670***** | | | 0.138 |
|  |  |  |  |  |  |  |  |  |  |  | Depth | | | **0.841***** | | | 0.162 |
|  |  |  |  |  |  |  |  |  |  |  | Wind exposure | | | **- 1.209***** | | | 0.169 |
|  |  | Depth * Wind | | - 229.2 | | 2.4 | | 8 | 0.230 | | Intercept | | | **- 2.701***** | | | 0.154 |
|  |  |  |  |  |  |  |  |  |  |  | Depth | | | **0.896***** | | | 0.200 |
|  |  |  |  |  |  |  |  |  |  |  | Wind exposure | | | **- 1.113***** | | | 0.262 |
|  |  |  |  |  |  |  |  |  |  |  | Depth*Wind exposure | | | - 0.1589 | | | 0.337 |
| CCA cover | | Depth * Wind | | - 137.5 | | 0.0 | | 8 | 0.833 | | Intercept | | | **- 3.134***** | | | 0.324 |
|  |  |  |  |  |  |  |  |  |  |  | Depth | | | 0.240 | | | 0.441 |
|  |  |  |  |  |  |  |  |  |  |  | Wind exposure | | | **1.870***** | | | 0.426 |
|  |  |  |  |  |  |  |  |  |  |  | Depth*Wind exposure | | | **- 1.672**** | | | 0.603 |
|  |  | Depth + Wind | | - 133.2 | | 4.2 | | 7 | 0.100 | | Intercept | | | **- 2.734***** | | | 0.301 |
|  |  |  |  |  |  |  |  |  |  |  | Depth | | | - 0.662 | | | 0.338 |
|  |  |  |  |  |  |  |  |  |  |  | Wind exposure | | | **1.087**** | | | 0.339 |
|  |  | Wind | | - 132.2 | | 5.2 | | 6 | 0.061 | | Intercept | | | **- 3.057***** | | | 0.278 |
|  |  |  |  |  |  |  |  |  |  |  | Wind exposure | | | **1.055**** | | | 0.363 |
| Fleshy macroalgae cover | | Depth | | - 183.8 | | 0.0 | | 6 | 0.700 | | Intercept | | | **- 4.353***** | | | 0.274 |
|  |  |  |  |  |  |  |  |  |  |  | Depth | | | **1.008**** | | | 0.336 |
|  |  | Depth + Wind | | - 181.4 | | 2.3 | | 7 | 0.220 | | Intercept | | | **- 4.287***** | | | 0.304 |
|  |  |  |  |  |  |  |  |  |  |  | Depth | | | **1.015**** | | | 0.335 |
|  |  |  |  |  |  |  |  |  |  |  | Wind exposure | | | - 0.156 | | | 0.335 |
|  |  | Depth * Wind | | - 179.1 | | 4.7 | | 8 | 0.067 | | Intercept | | | **- 4.370***** | | | 0.350 |
|  |  |  |  |  |  |  |  |  |  |  | Depth | | | **1.159**** | | | 0.446 |
|  |  |  |  |  |  |  |  |  |  |  | Wind exposure | | | 0.040 | | | 0.515 |
|  |  |  |  |  |  |  |  |  |  |  | Depth*Wind exposure | | | - 0.332 | | | 0.671 |
| Turf algae cover | | Depth * Wind | | - 97.2 | | 0.0 | | 7 | 0.523 | | Intercept | | | **- 1.143***** | | | 0.222 |
|  |  |  |  |  |  |  |  |  |  |  | Depth | | | - 0.086 | | | 0.292 |
|  |  |  |  |  |  |  |  |  |  |  | Wind exposure | | | 0.579 | | | 0.313 |
|  |  |  |  |  |  |  |  |  |  |  | Depth*Wind exposure | | | **0.893*** | | | 0.411 |
|  |  | Wind | | - 95.7 | | 1.5 | | 5 | 0.243 | | Intercept | | | **- 1.189***** | | | 0.173 |
|  |  |  |  |  |  |  |  |  |  |  | Wind exposure | | | 1.028 | | | 0.243 |
|  |  | Depth + Wind | | - 95.6 | | 1.6 | | 6 | 0.233 | | Intercept | | | **- 1.371***** | | | 0.204 |
|  |  |  |  |  |  |  |  |  |  |  | Depth | | | 0.367 | | | 0.233 |
|  |  |  |  |  |  |  |  |  |  |  | Wind exposure | | | **1.027***** | | | 0.236 |
|  |  | |  | |  | |  | | |  | |  | * P < 0.05 | | ** P < 0.01 | ***P < 0.001 | |

**Table S2**. AICc output and parameter estimates for the fixed effects of generalized linear mixed effects models examining the effects of depth and wind exposure on mean proportion of hard coral, CCA, fleshy macroalgae, and turf algae cover. Shallow and sheltered were set as baseline conditions for the model. Estimates significantly different from zero are bolded.

| Location | Shallow  mean cover | Deep  mean cover | Difference  (Deep – Shallow) |
| --- | --- | --- | --- |
| L7 | 0.0300 | 0.0378 | 0.0078 |
| L8 | 0.0117 | 0.0334 | 0.0217 |
| L9 | 0.0432 | 0.0660 | 0.0229 |
| L10 | 0.0062 | 0.0405 | 0.0343 |
| L11 | 0.0133 | 0.0611 | 0.0479 |
| L12 | 0.0410 | 0.0787 | 0.0376 |
| L13 | 0.0024 | 0.0163 | 0.0140 |
| L14 | 0.0085 | 0.0240 | 0.0156 |
| L15 | 0.0414 | 0.1116 | 0.0702 |
| L16 | 0.1365 | 0.0755 | - 0.0610 |
| L17 | 0.0536 | 0.1801 | 0.1265 |
| L18 | 0.0481 | 0.1524 | 0.1042 |
| L19 | 0.0537 | 0.1456 | 0.0919 |
| L20 | 0.0473 | 0.1163 | 0.0691 |
| L21 | 0.0765 | 0.1754 | 0.0989 |
| L22 | 0.0498 | 0.2302 | 0.1804 |

**Table S3**. Mean hard coral cover for deep and shallow sites at the 16 locations studied. Grey shading indicates sheltered locations.

| Taxon | Mean proportion cover | Relative coral cover (%) | Life History |
| --- | --- | --- | --- |
| *Porites* spp. | 0.0184 | 26.45 | Stress-tolerant |
| *Pavona duerdeni* | 0.0114 | 16.36 | Stress-tolerant |
| *Dipsastraea* spp. | 0.0043 | 6.26 | Stress-tolerant |
| *Platygyra* spp. | 0.0042 | 6.08 | Stress-tolerant |
| *Pavona varians* | 0.0039 | 5.6 | Stress-tolerant |
| Unknown Coral | 0.0036 | 5.24 | Unknown Coral |
| *Pocillopora* spp. | 0.0032 | 4.67 | Competitive |
| *Goniastrea stelligera* | 0.0026 | 3.79 | Stress-tolerant |
| *Fungiidae* spp. | 0.0025 | 3.60 | Stress-tolerant |
| *Favites* spp. | 0.0023 | 3.29 | Stress-tolerant |
| *Astrea* spp. | 0.0023 | 3.27 | Stress-tolerant |
| *Acropora* spp. | 0.0021 | 3.02 | Competitive |
| *Leptastrea* spp. | 0.0020 | 2.94 | Weedy |
| *Leptoseris myceteroides* | 0.0013 | 1.80 | Stress-tolerant |
| *Turbinaria* spp. | 0.0011 | 1.64 | Competitive |
| *Hydnophora microconos* | 0.0009 | 1.37 | Stress-tolerant |
| *Montipora spp.* | 0.0009 | 1.30 | Competitive |
| *Astreopora* spp. | 0.0007 | 1.08 | Stress-tolerant |
| *Echinophyllia* spp. | 0.0004 | 0.62 | Stress-tolerant |
| *Lobophyllia* spp. | 0.0003 | 0.44 | Stress-tolerant |
| *Gardineroseris planulata* | 0.0002 | 0.32 | Stress-tolerant |
| *Oxypora* spp. | 0.0002 | 0.26 | Not Classified |
| *Lobophytum* spp. | 0.0002 | 0.23 | Soft Coral |
| *Sarcophyton* spp. | 0.0001 | 0.22 | Soft Coral |
| *Sinularia* spp. | 0.0001 | 0.11 | Soft Coral |
| *Goniastrea edwardsi* | 0.0000 | 0.05 | Stress-tolerant |

**Table S4.** Hard and soft coral cover by taxon pooled across both depth and wind exposure levels. Mean proportion cover is the average proportion contributed by each taxon to total benthic cover which includes coral and non-coral substrates. Relative coral cover is the average proportion contributed by each taxon relativized to total hard and soft coral cover. Life history classifications (following Darling et al. 2012) are included for each hard coral taxon.

**Table S5**. AICc output and parameter estimates for the fixed effects of generalized linear mixed effects models examining the effects of depth and wind exposure on *Acropora* and *Pocillopora* total colony counts. Shallow and sheltered were set as baseline conditions for the model. Estimates significantly different from zero are bolded.

| Response | | Model | | AICc | | ΔAICc | | df | *w_i_* | | Predictor | | | Estimate | | | SE |
| --- | --- | --- | --- | --- | --- | --- | --- | --- | --- | --- | --- | --- | --- | --- | --- | --- | --- |
| *Acropora* count | | Depth | | 216.7 | | 0.0 | | 5 | 0.686 | | Intercept | | | **0.788***** | | | 0.228 |
|  |  |  |  |  |  |  |  |  |  |  | Depth | | | **- 1.204***** | | | 0.301 |
|  |  | Depth + Wind | | 218.7 | | 2.1 | | 6 | 0.244 | | Intercept | | | **0.667*** | | | 0.300 |
|  |  |  |  |  |  |  |  |  |  |  | Depth | | | **- 1.200***** | | | 0.300 |
|  |  |  |  |  |  |  |  |  |  |  | Wind exposure | | | 0.244 | | | 0.391 |
|  |  | Depth * Wind | | 221.3 | | 4.6 | | 7 | 0.069 | | Intercept | | | **0.660*** | | | 0.316 |
|  |  |  |  |  |  |  |  |  |  |  | Depth | | | **- 1.177**** | | | 0.432 |
|  |  |  |  |  |  |  |  |  |  |  | Wind exposure | | | 0.259 | | | 0.436 |
|  |  |  |  |  |  |  |  |  |  |  | Depth*Wind exposure | | | - 0.045 | | | 0.599 |
| *Pocillopora* count | | Depth | | 256.8 | | 0.0 | | 5 | 0.596 | | Intercept | | | 0.179 | | | 0.286 |
|  |  |  |  |  |  |  |  |  |  |  | Depth | | | 0.660 | | | 0.377 |
|  |  | Depth + Wind | | 259.2 | | 2.4 | | 6 | 0.180 | | Intercept | | | 0.221 | | | 0.336 |
|  |  |  |  |  |  |  |  |  |  |  | Depth | | | 0.663 | | | 0.377 |
|  |  |  |  |  |  |  |  |  |  |  | Wind exposure | | | - 0.090 | | | 0.376 |
|  |  | Wind | | 259.7 | | 2.9 | | 5 | 0.140 | | Intercept | | | 0.543 | | | 0.284 |
|  |  |  |  |  |  |  |  |  |  |  | Wind exposure | | | - 0.078 | | | 0.396 |
|  |  | Depth * Wind | | 260.7 | | 3.9 | | 7 | 0.083 | | Intercept | | | 0.416 | | | 0.377 |
|  |  |  |  |  |  |  |  |  |  |  | Depth | | | 0.301 | | | 0.517 |
|  |  |  |  |  |  |  |  |  |  |  | Wind exposure | | | - 0.490 | | | 0.547 |
|  |  |  |  |  |  |  |  |  |  |  | Depth*Wind exposure | | | 0.743 | | | 0.744 |
|  |  | |  | |  | |  | | |  | |  | * P < 0.05 | | ** P < 0.01 | ***P < 0.001 | |

| Species | Location | Shallow  mean colony count | Deep  mean colony count | Difference  (Deep – Shallow) |
| --- | --- | --- | --- | --- |
| *Acropora* | L7 | 0.1875 | 0.0000 | - 0.1875 |
|  | L8 | 0.2500 | 0.1250 | - 0.1250 |
|  | L9 | 1.0000 | 0.0000 | - 1.0000 |
|  | L10 | 0.6875 | 0.4375 | - 0.2500 |
|  | L11 | 0.4375 | 0.0625 | - 0.3750 |
|  | L12 | 0.1875 | 0.1875 | 0.0000 |
|  | L13 | 0.0625 | 0.0000 | - 0.0625 |
|  | L14 | 0.1250 | 0.0000 | - 0.1250 |
|  | L15 | 0.1250 | 0.0625 | - 0.0625 |
|  | L16 | 0.6875 | 0.3125 | - 0.3750 |
|  | L17 | 0.1875 | 0.0000 | - 0.1875 |
|  | L18 | 0.1875 | 0.1250 | - 0.0625 |
|  | L19 | 0.1250 | 0.0625 | - 0.0625 |
|  | L20 | 0.3125 | 0.0625 | - 0.2500 |
|  | L21 | 0.3125 | 0.0000 | - 0.3125 |
|  | L22 | 0.1875 | 0.0625 | - 0.1250 |
|  | L7 | 0.4375 | 0.0000 | - 0.4375 |
| *Pocillopora* | L8 | 0.0625 | 1.8125 | - 1.7500 |
|  | L9 | 0.2500 | 0.7500 | 0.5000 |
|  | L10 | 0.0000 | 0.3125 | 0.3125 |
|  | L11 | 0.1875 | 0.1875 | 0.0000 |
|  | L12 | 0.2500 | 0.1250 | - 0.1250 |
|  | L13 | 0.0625 | 0.2500 | 0.1875 |
|  | L14 | 0.0000 | 0.6250 | 0.6250 |
|  | L15 | 0.1250 | 1.0000 | 0.8750 |
|  | L16 | 0.8750 | 0.8125 | - 0.0625 |
|  | L17 | 0.1250 | 0.1250 | 0.0000 |
|  | L18 | 0.0000 | 0.2500 | 0.2500 |
|  | L19 | 0.1875 | 0.0000 | - 0.1875 |
|  | L20 | 0.1250 | 0.1875 | 0.0625 |
|  | L21 | 0.3125 | 0.1250 | - 0.1875 |
|  | L22 | 0.3125 | 0.3750 | 0.0625 |

**Table S6**. Mean *Acropora* and *Pocillopora* colony counts for deep and shallow sites at the 16 locations studied. Grey shading indicates sheltered locations.

| Substrate | Wind Exposure | Depth | Mean ± SE |
| --- | --- | --- | --- |
| Hard coral | Exposed | Shallow | 0.020 ± 0.003 |
|  |  | Deep | 0.043 ± 0.005 |
|  | Sheltered | Shallow | 0.063 ± 0.006 |
|  |  | Deep | 0.148 ± 0.014 |
| CCA | Exposed | Shallow | 0.233 ± 0.021 |
|  |  | Deep | 0.061 ± 0.005 |
|  | Sheltered | Shallow | 0.047 ± 0.008 |
|  |  | Deep | 0.049 ± 0.006 |
| Macroalgae | Exposed | Shallow | 0.009 ± 0.003 |
|  |  | Deep | 0.039 ± 0.008 |
|  | Sheltered | Shallow | 0.012 ± 0.003 |
|  |  | Deep | 0.047 ± 0.006 |
| Turf algae | Exposed | Shallow | 0.381 ± 0.022 |
|  |  | Deep | 0.561 ± 0.014 |
|  | Sheltered | Shallow | 0.256 ± 0.015 |
|  |  | Deep | 0.235 ± 0.011 |

**Table S7**. Mean and SE of cover by substrate for each depth and wind exposure level.

**Table S8**. Mean CCA cover for deep and shallow sites at the 16 locations studied. Grey shading indicates sheltered locations.

| Location | Shallow  mean cover | Deep  mean cover | Difference  (Deep – Shallow) |
| --- | --- | --- | --- |
| L7 | 0.2127 | 0.0695 | - 0.1431 |
| L8 | 0.0154 | 0.0565 | 0.0411 |
| L9 | 0.3622 | 0.0682 | - 0.2940 |
| L10 | 0.3803 | 0.0786 | - 0.3016 |
| L11 | 0.2473 | 0.0897 | - 0.1576 |
| L12 | 0.1600 | 0.0356 | - 0.1244 |
| L13 | 0.0000 | 0.0058 | 0.0058 |
| L14 | 0.4885 | 0.0882 | - 0.4003 |
| L15 | 0.0047 | 0.0953 | 0.0906 |
| L16 | 0.0440 | 0.0000 | - 0.0440 |
| L17 | 0.1874 | 0.0316 | - 0.1558 |
| L18 | 0.0192 | 0.1074 | 0.0882 |
| L19 | 0.0036 | 0.0411 | 0.0375 |
| L20 | 0.0405 | 0.1026 | 0.0621 |
| L21 | 0.0421 | 0.0107 | - 0.0314 |
| L22 | 0.0334 | 0.0048 | - 0.0286 |

| Location | Shallow  mean cover | Deep  mean cover | Difference  (Deep – Shallow) |
| --- | --- | --- | --- |
| L7 | 0.0127 | 0.0471 | 0.0343 |
| L8 | 0.0000 | 0.0072 | 0.0072 |
| L9 | 0.0106 | 0.0047 | - 0.0058 |
| L10 | 0.0024 | 0.0060 | 0.0036 |
| L11 | 0.0407 | 0.1089 | 0.0682 |
| L12 | 0.0062 | 0.0178 | 0.0116 |
| L13 | 0.0000 | 0.0000 | 0.0000 |
| L14 | 0.0012 | 0.0999 | 0.0987 |
| L15 | 0.0035 | 0.0343 | 0.0307 |
| L16 | 0.0572 | 0.0037 | - 0.0535 |
| L17 | 0.0072 | 0.1003 | 0.0931 |
| L18 | 0.0049 | 0.0082 | 0.0033 |
| L19 | 0.0071 | 0.0590 | 0.0518 |
| L20 | 0.0024 | 0.0240 | 0.0216 |
| L21 | 0.0049 | 0.0390 | 0.0341 |
| L22 | 0.0107 | 0.1049 | 0.0942 |

**Table S9**. Mean fleshy macroalgae cover for deep and shallow sites at the 16 locations studied. Grey shading indicates sheltered locations.

**Table S10**. Mean turf algae cover for deep and shallow sites at the 16 locations studied. Grey shading indicates sheltered locations.

| Location | Shallow  mean cover | Deep  mean cover | Difference  (Deep – Shallow) |
| --- | --- | --- | --- |
| L7 | 0.3462 | 0.5413 | 0.1951 |
| L8 | 0.6737 | 0.4622 | - 0.2114 |
| L9 | 0.2526 | 0.5011 | 0.2485 |
| L10 | 0.3733 | 0.5221 | 0.1488 |
| L11 | 0.1524 | 0.5607 | 0.4084 |
| L12 | 0.5156 | 0.7102 | 0.1946 |
| L13 | 0.6006 | 0.6892 | 0.0886 |
| L14 | 0.1355 | 0.6250 | 0.3375 |
| L15 | 0.1633 | 0.1485 | - 0.0148 |
| L16 | 0.2746 | 0.3765 | 0.1020 |
| L17 | 0.3014 | 0.2495 | - 0.0519 |
| L18 | 0.1247 | 0.1422 | 0.0175 |
| L19 | 0.1269 | 0.2787 | 0.1518 |
| L20 | 0.2552 | 0.2596 | 0.0044 |
| L21 | 0.2397 | 0.2866 | 0.0469 |
| L22 | 0.5630 | 0.1407 | - 0.4223 |

**Table S11**. Mean coral cover before and after the 2015-2016 El Niño caused a coral mass mortality event at very low disturbance (VL) locations on Kiritimati. Only shallow (10-12 m) sites were surveyed. Data from Baum et al. 2023. For a map of VL locations, see Figure S5.

| **Location** | **Wind Exposure** | **Before** | **After** |
| --- | --- | --- | --- |
| VL1 | Exposed | 60.692 | 3.444 |
| VL2 | Exposed | 58.897 | 1.097 |
| VL3 | Sheltered | 57.002 | 5.191 |
| VL5 | Exposed | 32.750 | 1.720 |
